# libRoadRunner: A High Performance SBML Compliant Simulator

**DOI:** 10.1101/001230

**Authors:** E. T. Somogyi, M. T. Karlsson, M. Swat, M. Galdzicki, H. M Sauro

## Abstract

**Summary:** We describe libRoadRunner, a cross-platform, open-source, high performance C++ library for running and analyzing SBML-compliant models. libRoadRunner was created primarily to achieve high performance, ease of use, portability and an extensible architecture. libRoadRunner includes a comprehensive API, Plugin support, Python scripting and additional functionality such as stoichiometric and metabolic control analysis.

**Accessibility and Implementation:** To maximize collaboration, we made libRoadRunner open source and released it under the Apache License, Version 2.0. To facilitate reuse, we have developed comprehensive Python bindings using SWIG (swig.org) and a C API. LibRoadRunner uses a number of statically linked third party libraries including: LLVM [4], libSBML [1], CVODE, NLEQ2, LAPACK and Poco. LibRoadRunner is supported on Windows, Mac OS X and Linux.

**Supplementary information:** Online documentation, build instructions and git source repository are available at http://www.libroadrunner.org

## 1 Introduction

Over last few decades researchers have developed many tools to simulate biochemical networks [7]. Most of such simulators are however embedded in monolithic environments, making it very difficult to reuse the code in other applications. To resolve this issue various groups have built simulators as reusable libraries. Most notable of these are COPASI [2], libSBMLSim [9], RoadRunner C# [7], SOSLib [5] and SBSCL [3]. Despite all these efforts, finding a simulator that is easy-to-use, has good SBML-compliance, is capable of handling complex models and fast enough to run thousands of models is challenging; for example, in multi-cell simulators such as CompuCell3D. We designed libRoadRunner to address these shortcomings and by using Just-In-Time (JIT) compilation via Low Level Virtual Machine (LLVM), we have significantly improved computational performance as compared to existing solutions.

This version of libRoadRunner has a new internal design compared with the previously developed C# roadRunner. The APIs are designed with the modeler in mind, offer a wide range of functionality and provide high performance access to model and simulation data, making libRoadRunner ideal for realtime interactive environments. To facilitate easy extensibility libRoadRunner provides a plugin API making it possible to easily write extensions. Figure 1 shows the architectural overview of libRoadRunner.

The most novel and important component of the libRoadRunner is its computational core based on the LLVM. LLVM is designed for real-time optimization and dynamic compilation of computer languages. This has allowed us to achieve extremely low runtimes. This speed is particularly important for large simulation applications such as multi-cell models that require many thousands of biochemical networks to be simulated simultaneously. libRoadRunner 1.0 supports all models in the SBML test suite except for those that include delay equations and algebraic rules.

## 2 Methods

### 2.1 LLVM Backend

We have developed a non-tracing, multi-pass JIT compiler for the SBML language using the LLVM framework which allows us to directly JIT-compile SBML in memory into native machine code at runtime. Our compiler treats SBML as a statically typed, dynamically scoped declarative language allowing us to not only generate specific code for each SBML model, but to also generate a specific data layout that is optimized for each model. As a result the model and the integrator share the same data layout eliminating the need to copy data between the integrator and the model evaluation code. The flexible architecture of the compiler also greatly facilitates additions of SBML extensions

### 2.2 Extensibility

A key design requirement of libRoadRunner was to provide a component-based architecture. Therefore, internal components such as the integrator, steady state solver, SBML compiler, etc., only communicate with each other through pure virtual interfaces. This design allows us to readily add new internal components such as a planned GPU SBML compiler, new integrators, stochastic integrators and steady state solvers. While the easiest way to extend functionality is to write Python scripts using the Python API, for performance sensitive applications developers can access the same functionality through the C++ or C API’s. We also offer a C Plugin API that allows developers to write dynamically loadable plugins in any C complaint language such as Fortran or Object Pascal. The Plugin API is written such that it allows generic access to all plugins. For example, scripting languages such as Python can access a plugin without having to implement a new Python API for every plugin. The current libRoadRunner distribution comes with two example plugins that illustrate how libRoadRunner functionality can be extended. One example implements the Levenberg-Marquardt algorithm [6] for fitting models to experimental data.

### 2.3 Python API

LibRoadrunner provides a comprehensive and *pythonic* Python module which is designed to have the feel of the Python SciPy package (scipy.org) and exposes the full functionality of the public C++ API. Consequently, the easiest way to use libRoadRunner is via Python scripting. This API allows access to any SBML element along with calculated values such as the Jacobian, eigenvalues or the various control coefficients using the Python dictionary protocol. All libRoadRunner data arrays are accessed directly without copying as Numpy arrays. This provides a commonly used access protocol. Using the Python API, modelers can create one or many RoadRunner simulator objects which offers a very natural way of using libRoadRunner as the Reaction-Kinetics engine in multi-cell simulators such as CompuCell3D (www.compucell3d.org).

### 2.4 Performance

To demonstrate the capabilities of libRoadRuner we compared it to three simulator libraries: libSBMLSim, COPASI and SBSCL. We tested the ability to pass the SBML test suite (1008 models), the ability to simulate very large models and the ability to simulate systems with large numbers of events. Table 1 lists the times recorded from four different libraries based on three types of models. The first model tests the ability to scale to large numbers of state variables defined by relatively simple ODEs. The second model tests the ability to deal with complex SBML functions. The timings are the total wall time to complete each process. Tests were run on a 2.6 GHz Mac Pro, OS X 10.6, and used COPASI v. 4.9.43, LibSMLSim v. 1.1 and SBSCL v. 1.3. Note, we experienced instability of LibSBMLSim runs with *>*100 copies of the Brusselator system or complex SBML.

**Figure 1.**
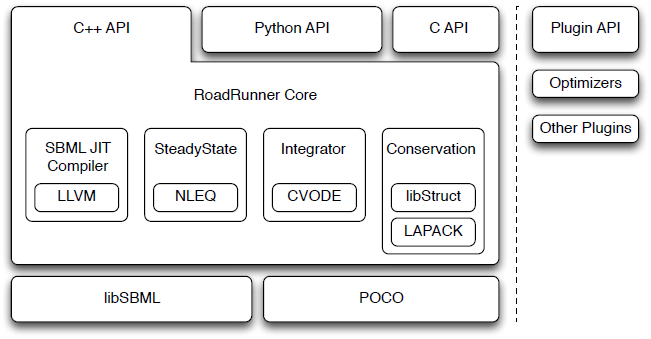
Architectural Overview of libRoadRunner.

### 2.5 Using libRoadRunner in multi-cell simulations

We have successfully deployed libRoadRunner as the Reaction-Kinetic solver engine in CompuCell3D - a simulation environment used to build multi-cell based models of tissues. We have used the libRoadRunner Python API to associate each cell with a collection of libRoadRunner-based simulators. Since CompuCell3D uses Python scripting to describe its models, integrating libRoadRunner was straightforward.

**Table 1.**
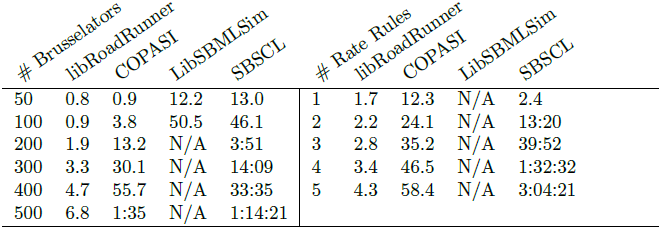
All times are in seconds. The first set of tests consisted of *N* copies of the Brusselator system in a single model. The second set was a sin function implemented as a 63 element piecewise SBML functions combined with *N* parameters defined by rate rules integrating the sin function. Test models are available at libroadrunner.org.

| # Brusselators | libRoadRunner | COPASI | LibSBMLSim | SBSCl | # Rate Rules | libRoadRunner | COPASI | LibSBMLSim | SBSCl |
| --- | --- | --- | --- | --- | --- | --- | --- | --- | --- |
| 50 | 0.8 | 0.9 | 12.2 | 13.0 | 1 | 1.7 | 12.3 | N/A | 2.4 |
| 100 | 0.9 | 3.8 | 50.5 | 46.1 | 2 | 2.2 | 24.1 | N/A | 13:20 |
| 200 | 1.9 | 13.2 | N/A | 3:51 | 3 | 2.8 | 35.2 | N/A | 39:52 |
| 300 | 3.3 | 30.1 | N/A | 14:09 | 4 | 3.4 | 46.5 | N/A | 1:32:32 |
| 400 | 4.7 | 55.7 | N/A | 33:35 | 5 | 4.3 | 58.4 | N/A | 3:04:21 |
| 500 | 6.8 | 1:35 | N/A | 1:14:21 |  |  |  |  |  |

We ran the Delta-Notch patterning simulation [8] on an array of 36 adjacent cells with different reaction-kinetics solvers and determined that the libRoad-Runner solver led to a two fold improvement in runtime. In multi-cell simulations time spent on solving reaction-kinetics models makes up only a portion of the total runtime. Thus the two fold speedup in the overall simulation time is significant.

## 3 Discussion

Decades of development has produced powerful, intuitive and feature-rich software to simulate biochemical networks. The existing portfolio of SBML-compliant tools satisfies the needs of most modelers. Yet, limitations of existing packages present a significant research barrier for applications that require high performance, excellent SBML compatibility, broad functionality and extensibility. LibRoadRunner addresses these shortcomings. It is designed for researchers who need a simulation engine for interactive desktop simulation, multicellular simulations, which often require 10,000 or more simultaneous simulations, and for web based interactive simulation with multi-session requirements. Even though the current version of the RoadRunner appears to be complete, we are planning a number of additional features including even better SBML compliance, stochastic GPU integrators, parallel distributed model JIT compilation and simulation, spatial models as well as new plugins such as alternative optimizers, and bifurcation analysis. By using an open-source, open-access development process we hope to engage external developers which should result in higher quality and more frequent future releases of libRoadRunner.

## Acknowledgment

We wish to acknowledge Frank Bergmann who with Herbert Sauro wrote the original roadRunner library in C# and wrote libStruct, the structural analysis library along with Ravi Rao. Finally we wish to thank Stanley Gu for testing libRoadRunner in his real time web based environment. **Funding:** Work supported by the National Institute of General Medical Sciences of the National Institutes of Health under award numbers R01-GM081070. The content is solely the responsibility of the authors and does not necessarily represent the official views of the National Institutes of Health.

